# Simultaneous mesoscopic measurement and manipulation of mouse cortical activity

**DOI:** 10.1101/2024.11.01.621418

**Authors:** Pascha Matveev, Anna J. Li, Zhiwen Ye, Anna J. Bowen, Ximena Opitz-Araya, Jonathan T. Ting, Nicholas A. Steinmetz

## Abstract

Dynamics of activity across the cerebral cortex at the mesoscopic scale — coordinated fluctuations of local populations of neurons — are essential to perception and cognition and relevant to computations like sensorimotor integration and goal-directed task engagement. However, understanding direct causal links between population dynamics and behavior requires the ability to manipulate mesoscale activity and observe the effect of manipulation across multiple brain regions simultaneously. Here, we develop a novel system enabling simultaneous recording and manipulation of activity across the dorsal cortex of awake mice, compatible with large-scale electrophysiology from any region across the brain. Transgenic mice expressing the GCaMP calcium sensor are injected systemically with an adeno-associated virus driving expression of the ChrimsonR excitatory opsin. This strategy drives expression of the blue-excited calcium indicator, GCaMP, in excitatory neurons and red-excited Chrimson opsin in inhibitory neurons. We demonstrate widefield single-photon calcium imaging and simultaneous galvo-targeted laser stimulation over the entire dorsal cortical surface. The light channels of the imaging and the opsin do not interfere. We characterize the spatial and temporal resolution of the method, which is suitable for targeting specific cortical regions and specific time windows in behavioral tasks. The preparation is stable over many months and thus well-suited for long-term behavioral experiments. This technique allows for studying the effect of cortical perturbations on cortex-wide activity, on subcortical spiking activity, and on behavior, and for designing systems to control cortical activity in closed-loop.

## Introduction

A central challenge in systems neuroscience is causally linking behavior to neural activity patterns in defined brain regions or cell populations. For decades, lesions and pharmacological manipulations were the key techniques for establishing causal contributions of brain regions but these have significant limitations in their slow timecourses and imprecise targeting. Optogenetics now provides a method for fast-timescale, spatially precise, and cell-type specific manipulations. But linking effects of manipulations directly to behavioral outcomes, while powerful, has proven to be misleading in some cases (Otchy et al., 2015). Indeed, any manipulation likely produces diverse and far-ranging effects on activity around the brain, effects that may vary from trial-to-trial. The ability to simultaneously read and perturb activity patterns could overcome this challenge in two ways. First, we could directly observe the outcome of the manipulation on each trial, and thus link the specific stimulus-induced changes of brain activity to the behavioral outcomes. Second, we could in principle design closed-loop control algorithms to reliably achieve the desired manipulation pattern on each trial.

“Widefield” imaging, which uses a low-magnification, large field-of-view microscope and single-photon imaging of intrinsic signals or fluorescent reporters, has been used for decades to measure neural activity across large areas of cortex (Orbach et al., 1985; Grinvald et al., 1986). In recent years,application of this approach to mice — whose thin skulls permit transcranial imaging, and with transgenic brain-wide expression of high signal-to-noise calcium sensors — has yielded high spatial and temporal resolution of imaging across all dorsal cortical areas (Vanni and Murphy, 2014; Wekselblatt et al., 2016). This mesoscale imaging approach has been used to study the relationship of cortical dynamics to perception, action, and cognition (Allen et al., 2017; Makino et al., 2017; Musall et al., 2019; Couto et al., 2021; Zatka-Haas et al., 2021). In parallel, the same characteristics of mice (transparent skulls and transgenic suitability) have enabled the application of optogenetics across cortical areas (Guo et al., 2014; Pinto et al., 2019; Zatka-Haas et al., 2021). These experiments typically use a similar preparation, expressing an excitatory opsin in inhibitory neurons to suppress cortical activity around the stimulated region (Atallah et al., 2012; Li et al., 2019). Moreover, it is possible to combine electrophysiology with both widefield imaging (Xiao et al., 2017; Clancy et al., 2019; Peters et al., 2021; Ye et al., 2023) and with transcranial optogenetics (Zatka-Haas et al., 2021). Therefore, we hypothesized that with a suitable genetic strategy and microscope design, it would be possible to combine all three techniques in a single experiment.

Here we introduce a microscope design and genetic strategy for simultaneously measuring and manipulating brain activity across the whole dorsal cortical surface in awake, head-fixed mice using single photon widefield calcium imaging and optogenetics. We use transgenic GCaMP6s (Wekselblatt et al., 2016) or 8s (Wang et al., 2023) mice retro-orbitally injected with a Chrimson virus using the PHP-eB serotype (Challis et al., 2019). The Chrimson opsin is expressed in inhibitory neurons of the forebrain via the DLX2.0 promoter (Lee et al., 2023), and the GCaMP is expressed in excitatory neurons (CaMKII driver). We show that precise, graded measurement and manipulation are simultaneously possible with this strategy, and the measurement and manipulation spectra do not interfere. Using these techniques in tandem with electrophysiological recordings, we demonstrate a three-modality experiment with whole-brain and multi-timescale access. We use this combination to characterize the relationship between underlying single-neuron spiking activity and the imaging and stimulation paradigm. Additionally, this technique can be used many times within a single subject with repeatable results over at least 4 weeks.

Our approach serves as a way to measure the spatiotemporal effect of cortical perturbations on large-scale neural dynamics.

## Results

### An optical and genetic strategy for mesoscale measurement and manipulation

We designed a custom microscope for widefield calcium imaging across cortical regions (Vanni and Murphy, 2014; Couto et al., 2021; Zatka-Haas et al., 2021) that combines light paths for calcium fluorescence imaging with a galvanometer-directed laser beam (Figure 1A, Table 2). Frames were alternately illuminated with either blue (470 nm) or violet (405 nm) light, enabling correction for signal changes related to blood flow (Kim et al., 2016; Zatka-Haas et al., 2021). A 638 nm laser was fixed above a scan and tube lens pair and controlled by two mirror galvanometers. This configuration achieves imaging at a resolution of 17.3 µm/pixel and 70 frames per second (35 frames per second after hemodynamic correction, see Methods) with a field of view of up to 14.2 × 11.8 mm (820 × 685 pixels after 3 × 3 binning, Supplemental Figure 1A). The laser spot (0.12 mm diameter, Supplemental Figure 1C, see Methods) can be directed to any location on the dorsal cortical surface (Supplemental Figure 1B). The working distance of the microscope is 7.5 cm, leaving sufficient space to insert recording electrodes between the objective lens and the skull.

**Figure 1.**
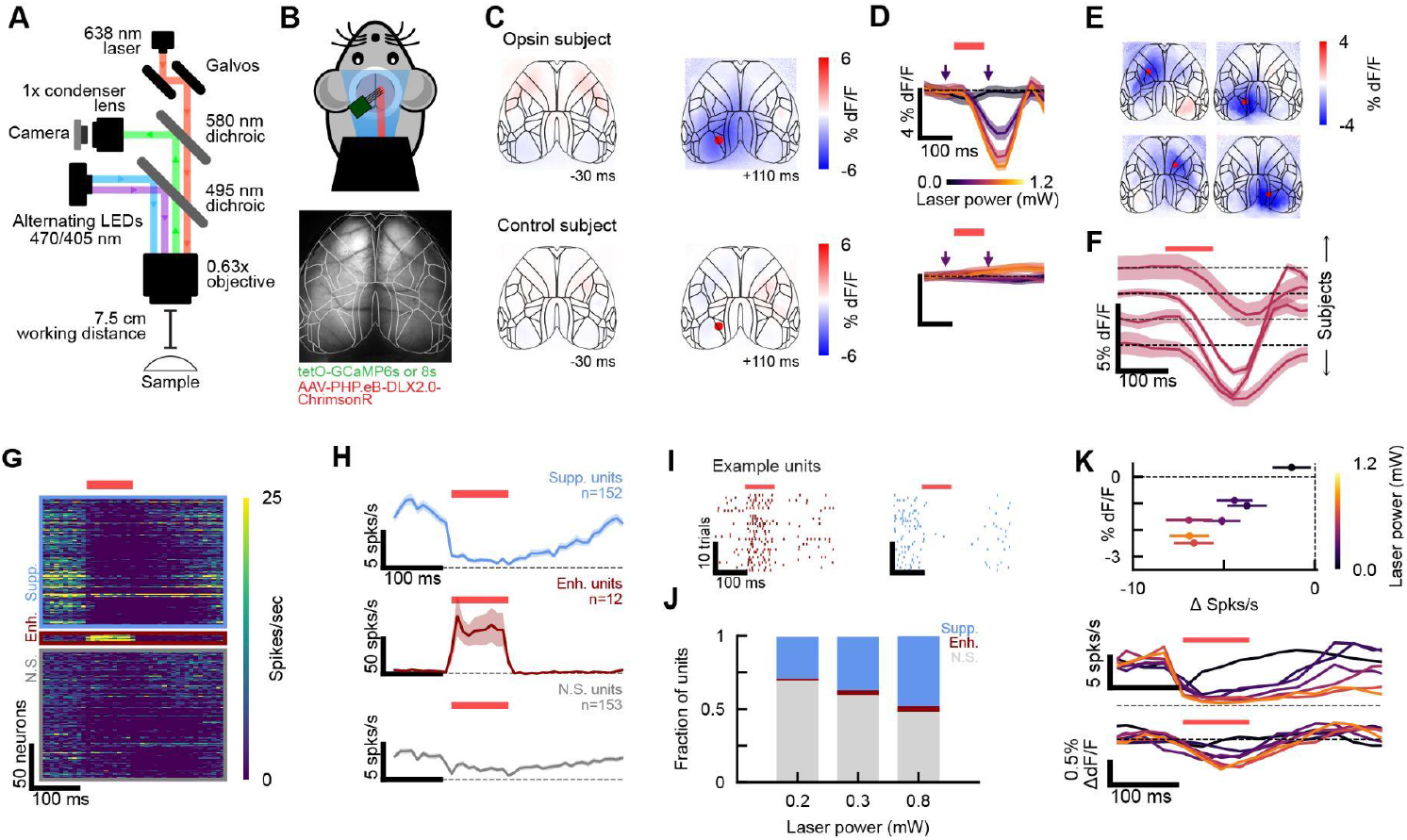
An imaging and genetic strategy for simultaneous measurement and manipulation of mesoscopic cortical activity in mice. A. Left, schematic of the microscope optical path. Blue (470 nm) and violet (405 nm) LEDs reflect off the first dichroic (495 nm cutoff) toward the skull. Emitted green photons pass through the first dichroic and reflect off the second dichroic (580 nm cutoff) to the camera. A red (638 nm) laser is steered with a two-mirror galvanometer and passes through both dichroics to the brain. Not to scale. B. Top, schematic of mouse experimental setup. Bottom, example mean frame from widefield imaging. C. Top row, mean dF/F images of calcium response to laser stimulation (n=45 trials). Left, 30 ms before onset. Right, 110 ms after onset. Bottom, as top but for “control” subject expressing GCaMP but no opsin (n=30 trials). Red dot indicates laser target. D. Top, timecourse of calcium response (solid line, mean; shaded regions; S.E.M. across trials) across laser powers. Red bar indicates laser on. Arrows indicate trial times for example images in C. Bottom, same as top but for control subject. E. Mean dF/F images of calcium response to different laser target locations. Images are 130 ms after stimulus onset. Red dot indicates laser target. F. Calcium response to 0.6 mW stimulus across subjects (bottom to top: Subject 1, n=50 trials; Subject 2, n=40 trials; Subject 3, n=40 trials, Subject 4, n=26 trials; example session per subject). Subject 4 (top) uses GCaMP6s compared to 8s in the other subjects (see Table 1) and accordingly has a slower response with lower signal-to-noise. G. Single unit responses (measured with electrophysiology) to laser stimulation (n=30 trials, example session). Units are categorized (colored boxes) by significance and direction of laser response (paired t-test between baseline and stimulus spike counts, p<0.05 per unit). H. Timecourse of laser responses averaged across neurons within categories from G. I. Left, trial-aligned raster for example opsin-expressing inhibitory neuron (from significantly enhanced category in G-H). Right, trial-aligned raster for example suppressed neuron (from significantly suppressed category in G-H). J. Fraction of units in each response category across laser powers. K. Top, magnitude of inactivation as a function of laser power measured in widefield imaging and electrophysiology. Error bars are S.E.M. across neurons. Middle, timecourse of inactivation measured in electrophysiology. Bottom, as middle for widefield imaging.

As our optical strategy combines imaging of green fluorescence with red-wavelength stimulation, we designed a genetic strategy for cortex-wide expression of GCaMP alongside the red-shifted channelrhodopsin Chrimson (Klapoetke et al., 2014). While in principle this strategy could be reversed, i.e., a red-shifted calcium sensor (Dana et al., 2016) and a blue-shifted opsin (Boyden et al., 2005; Klapoetke et al., 2014), the signal-to-noise ratio (SNR) for red-shifted indicators is significantly lower than for GCaMP, and fewer transgenic mouse lines are available. A potential disadvantage of our strategy is that red-shifted opsins have weak short-wavelength sensitivity (Klapoetke et al., 2014), but this does not create a confound in practice (Supplemental Figure 2B). To enable cortical manipulations at any location, we aimed to achieve even expression of the opsin transgene by using an AAV PHP.eB serotype virus, injected retro-orbitally during development (between P28-42). The opsin is expressed in forebrain GABAergic neurons under the DLX2.0 promoter (Lee et al., 2023).

**Figure 2.**
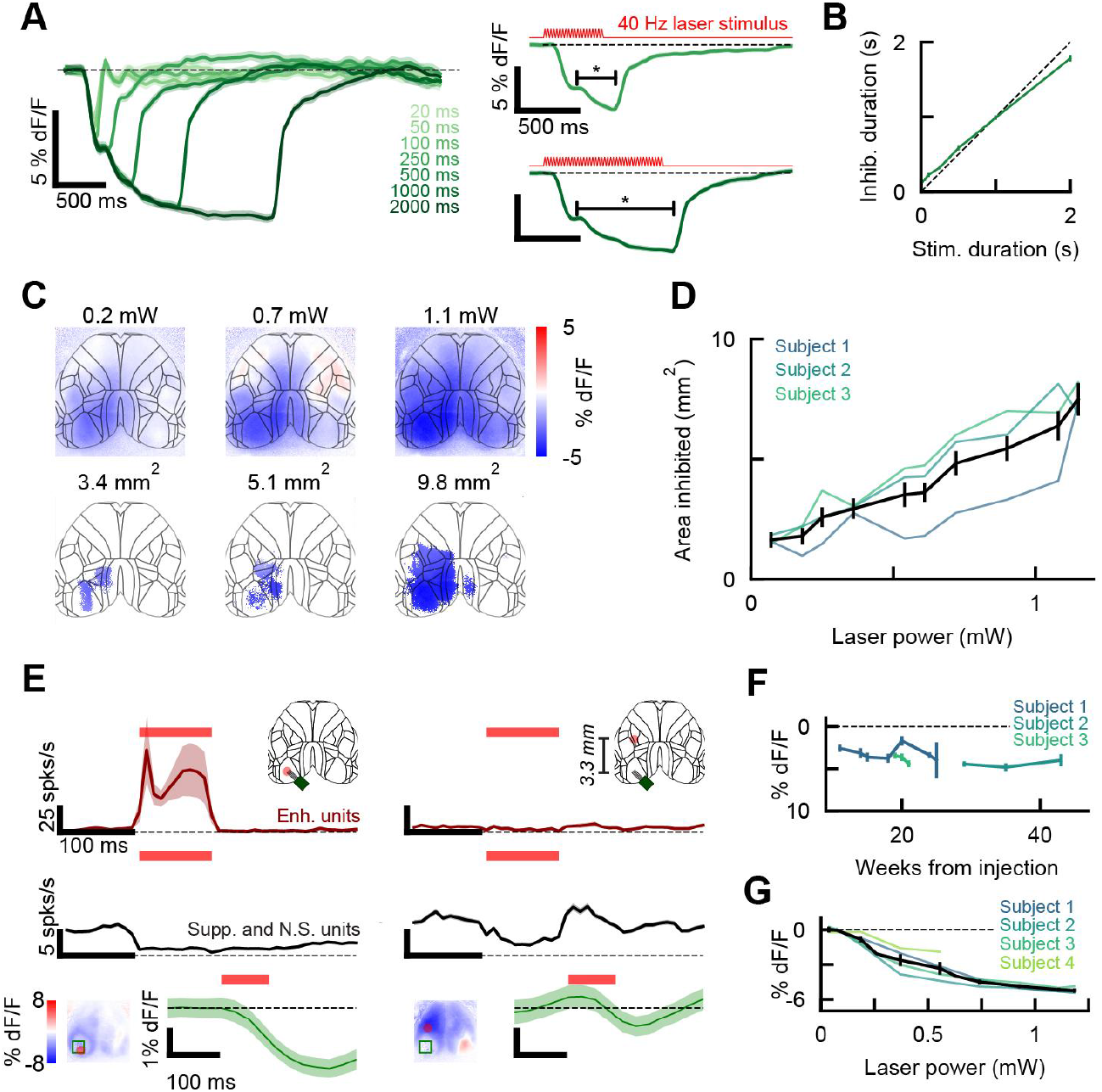
Temporal and spatial resolution of optogenetic manipulation. A. Left, calcium responses to different stimulus durations for an example session (solid line, mean; shaded region, S.E.M. across trials; n=35 trials, 0.7 mW laser power). Right, two example duration conditions. Red line indicates laser stimulus shape and duration. Bracket indicates time of significant suppression (p<0.05 based on a null distribution of baseline activity, see Methods). B. Inactivation duration (observed in widefield, significance test as in A) as a function of stimulus duration (n=2 subjects, n=180 trials per condition per subject, 0.7 mW laser power). C. Top, example images of calcium activity 150 ms after laser onset in response to different laser powers (example session, n=30 trials per condition). Bottom, masked images showing significantly inactivated area (p<0.03 from null distribution of baseline activity, see Methods). D. Average cortical inactivation area as a function of laser power across subjects (n=120 trials per condition per subject). Black line indicates mean and S.E.M. across all trials. E. Laser stimulation effects, measured with electrophysiology, at 3.3 mm from laser target. Top row, spiking activity of opsin-expressing neurons (n=12, same as in Fig. 1H). Schematic indicates laser target (red dot) and probe location (green square). Middle row, spiking activity of all other recorded neurons (n=305). Bottom row, mean calcium activity from green ROI indicated in image (n=30 trials). F. Calcium activity (130 ms after laser onset) for each subject as a function of weeks from injection (0.6 mW laser power). G. Calcium activity (130 ms after laser onset) as a function of laser power across subjects (Subject 1, n=7 sessions; Subject 2, n=3 sessions; Subject 3, n=3 sessions, Subject 4, n=4 sessions). Black line indicates mean and SEM.

### Simultaneous imaging and optogenetic stimulation

We achieved reliable simultaneous calcium imaging and optogenetic manipulation in all four opsin-expressing mice (Figure 1F, Table 1). Suppression at the target location occurred within one frame (∼29 ms, Figure 1D). We observed consistent inhibitory responses at multiple target laser locations on the dorsal cortical surface (Figure 1E). In all subjects, the minimum detectable response to laser stimulation occurred at 0.18 mW across the power levels tested (n=4, Supplemental Figure 2A).

**Table 1.**

| Mouse ID | Sex | Genotype | Injected virus | Water restricted for sessions | WF sessions | WF +ephys sessions | Figure panels |
| --- | --- | --- | --- | --- | --- | --- | --- |
| AL_0033<br>“Subject 1” | M | tetO-GCaMP 8s | AAV-PHP.eB-DLX 2.0-ChrimsonR-td Tomato, 1.6e12 GC in 100 uL PBS | Yes | 10 | 0 | 1F, 2B, 2D, 2G |
| AB_0032<br>“Subject 2” | F | tetO-GCaMP 8s | AAV-PHP.eB-DLX 2.0-ChrimsonR-td Tomato, 1.6e12 GC in 100 uL PBS | Yes | 4 | 2 | 1B-K, 2C-G |
| AL_0034<br>“Subject 3” | F | tetO-GCaMP 8s | AAV-PHP.eB-DLX 2.0-ChrimsonR-td Tomato, 1.6e12 GC in 100 uL PBS | Yes | 6 | 1 | 1F, 2A-B, 2D, 2F-G |
| ZYE_0069<br>“Subject 4” | F | tetO-GCaMP 6s | AAV-PHP.eB-DLX 2.0-ChrimsonR-td Tomato, 2.7e12 GC in 150 uL PBS | No | 2 | 0 | 1F, 2G |
| AL_0035<br>“Subject 5” | F | tetO-GCaMP<br>8s | none | Yes | 1 | 0 | 1C, 3A-B,<br>D, F-J |
| AL_0023<br>“Subject 6” | M | tetO-GCaMP<br>8s | none | No | 1 | 0 | 3D, G-H, J |
| ZYE_0087<br>“Subject 7” | F | tetO-GCaMP<br>8s | none | Yes | 1 | 0 | 3D, G-H, J |
| ZYE_0088<br>“Subject 8” | F | tetO-GCaMP<br>8s | none | Yes | 1 | 0 | 3D, G-H, J |
| AL_0023<br>“Subject 9” | M | tetO-GCaMP<br>8s | none | Yes | 1 | 0 | 3C |

**Table 2.**

| **Item** | **Manufacturer** | **Part number** |
| --- | --- | --- |
| 60 mm cage system for optics | Thorlabs | ER series rods, LCP09 cage plates |
| Laser | Omicron Laserage | LuxX 638nm-200mW |
| Laser fiber | Thorlabs | M42L01 |
| Collimating lens | Thorlabs | F280FC-A |
| Galvos | Thorlabs | GVS002 |
| Galvo/laser mount | Thorlabs | GCM102/M |
| Laser to mount coupler | Thorlabs | AD11F |
| Lens tube | Thorlabs | SM2V15 |
| Scan lens, $f = 100$ mm | Edmund Optics | 45-353 |
| ND filter | Thorlabs | NE2R09A |
| Tube lens, $f = 300$ mm | Thorlabs | AC508-300-AB |
| Camera | Basler | acA2440-75m |
| **Blue/violet imaging LEDs** |  |  |
| OptoLED power supply | Cairn | P1110/002/000 |
| Violet OptoLED head | Cairn | P1105/405/LED |
| Blue OptoLED head | Cairn | P1105/470/LED |
| Dual port microscope coupling | Cairn | P1210/CON/002 |
| Filter cube for emission filters | Cairn | P290/000/200 |
| Longpass dichroic beamsplitter (425 nm; emission filter and combiner for blue LED) | Chroma | T425lpxr-UF2 |
| Bandpass filter (395-415 nm; emission filter for violet LED) | Chroma | ET405/20x |
| UV-transmitting liquid light guide, 3 mm core diameter | Cairn | P135/015/003 |
| LED head to fiber optic coupler | Cairn | P1121/000/000 |
| **THT Macroscope (SciMedia)** |  |  |
| Epifluorescence box with 495 nm dichroic mirror | BrainVision | THT-FLSP |
| Epifluorescence box with 580 nm dichroic mirror and 530/55 nm emission filter | BrainVision | THT-FLSP |
| Condensing lens coupler for camera | BrainVision | CDAD-1 |
| Condensing lens | Leica | PlanApo 1x M-series |
| Objective | Leica | PlanApo 0.63x M-series |

Although Chrimson is weakly sensitive to short wavelength light, we found no response from opsin-expressing neurons when the blue and violet imaging LEDs were present without laser stimulation (Supplemental Figure 2B). This is likely due to the low light levels needed to obtain high signal-to-noise ratios in widefield imaging (0.012 mW/mm^2^ for blue; 0.019 mW/mm^2^ for violet, compared to 15.9 mW/mm^2^ for laser stimulation at the power for minimum detectable response, see Methods).

A 580 nm longpass dichroic placed before the imaging camera prevented interference from the red laser. In a control mouse without the Chrimson opsin, there were no excitatory or inhibitory effects observed in the widefield imaging due to laser stimuli (Figure 1C, D).

### Electrophysiological recordings

The effects of opsin activation observed in widefield calcium imaging were consistent with data from electrophysiological recordings. We used a Neuropixels 2.0 probe to record from cortical neurons (n=3 sessions, n=2 subjects, n=1019 total neurons, 339±11 neurons per session). To minimize light artifacts of both imaging and stimulation light sources, we designed each to have ramped onsets and offsets over short periods (see Methods). We identified an average of 3.1±0.7% opsin-expressing inhibitory neurons, which responded within 5 ms of laser onset (Figure 1G-I). These neurons cannot be observed in widefield imaging as the GCaMP expression is restricted to excitatory neurons. Suppression of activity in other neurons began within 10.6±0.5 ms, and lasted up to 129±3 ms (100 ms laser stimulus; Figure 1H, I). At sufficiently high power (0.8 mW), the laser achieved 62±7% suppression of the local spiking activity (Figure 1H), with significant suppression of 48% of recorded non-optotagged neurons (Figure 1J, paired t-test between activity 100 ms before and after stimulus onset, n=30 trials). The magnitude of inhibition as a function of laser power was strongly correlated between electrophysiology and widefield imaging (Figure 1K, Pearson’s r, r= 0.93, p=0.002).

### Temporal and spatial properties of inactivation

Different parameters of the laser stimulus elicited different patterns of inactivation.We reliably controlled the duration of cortical inactivation using a 40 Hz sinusoidal stimulus, which reduced adaptation of the photosensitive ion channels (Li et al., 2019) (Figure 2A). The period of inactivation was linearly related to the duration of laser stimulus, with a slope slightly below unity (slope=0.83, intercept=0.13, Figure 2B). We also observed that higher laser powers produced a larger area of inactivation (Figure 2C, D). This effect is mediated by stronger synaptic effects at higher laser powers (rather than increased laser scattering, for example). Consistent with widefield imaging results, inhibition was observed in electrophysiology at 3.3 mm away from the laser target (Figure 2E).

At this distance, no opsin-expressing neurons exhibited enhanced firing rates, suggesting that direct opsin activation is more localized than the range of inhibition (Figure 2E).

### Stability across time

Within subjects, the effect of optogenetic stimulation was consistent over many weeks (Figure 2F), with stable inactivation magnitudes observed up to 43 weeks after injection. In the first subject (Subject 4), we could no longer drive the opsin at 52 weeks after injection, though we did not have standardized quantitative data to demonstrate this. In subsequent subjects, we implemented a repeated test over weeks. There was no quantitative decline in opsin performance for these subjects. The monotonic relationship between power and inactivation magnitude was consistent across subjects (Figure 2G, Supplemental Figure 2A).

### Retinal photoreceptor response to laser stimulation

Given the observation that longer-wavelength light can penetrate more deeply into tissue, we investigated the possibility that our red laser stimuli might activate retinal photoreceptors and induce an unintended visual response. To test potential visual effects without interference from optical inhibition, we performed laser stimulus experiments in transgenic GCaMP8s mice with no viral injection for opsin expression (Table 1). We indeed observed increased activity in primary visual cortex (VISp) when stimulating frontal cortex, which is near the retina (Figure 3A). This visual response was dependent on laser power; at 0.9 mW, the visual response was comparable in magnitude to a 12.5% contrast full-field checkerboard visual stimulus (Figure 3C, B). We did not observe a significant visual response when stimulating somatosensory or visual areas of cortex (Figure 3D, Supplemental Figure 3B).

**Figure 3:**
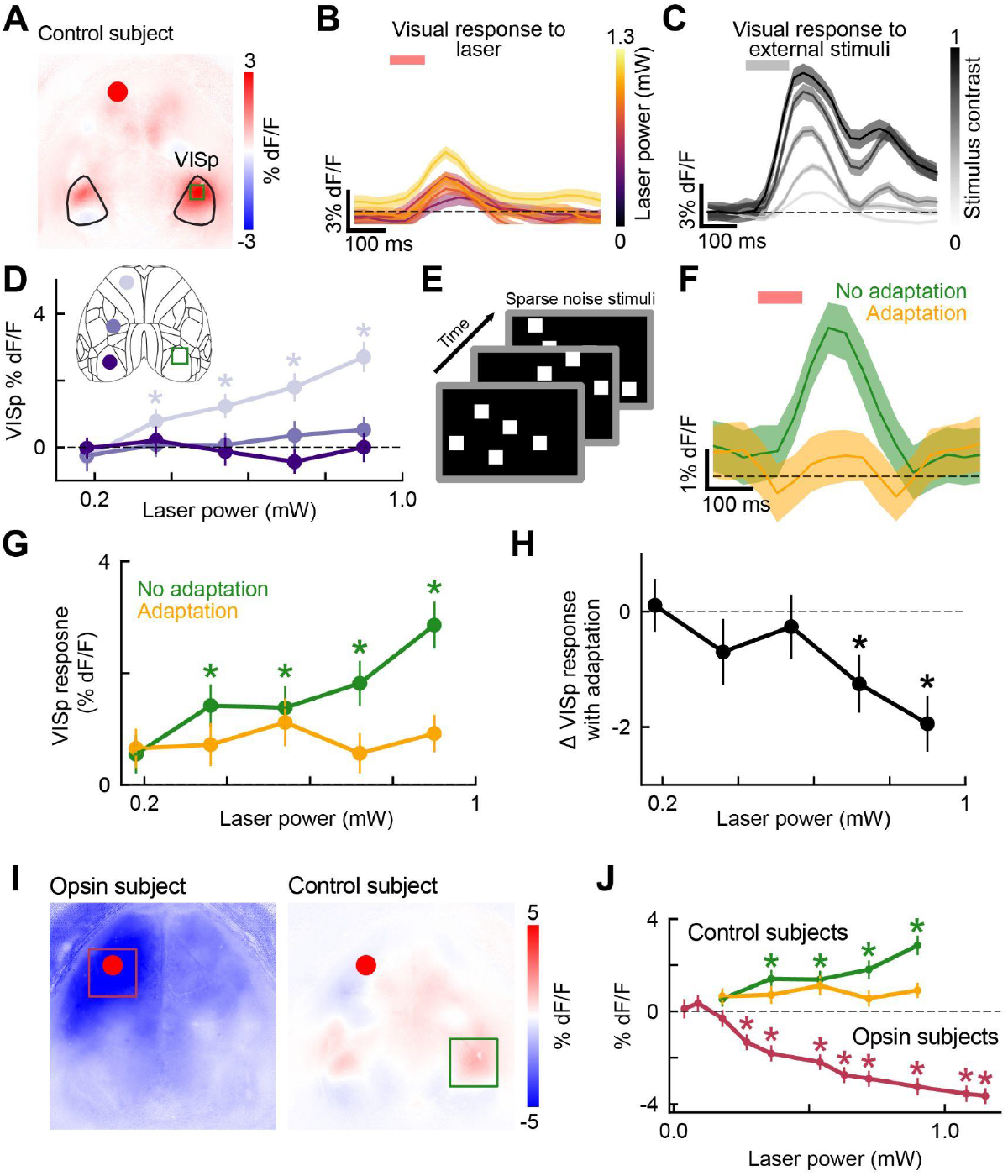
Excitatory responses caused by direct retinal activation in the visual cortex can be adapted. A. Mean image of off-target VISp response 150 ms after laser onset (example session, n=35 trials). Red dot indicates laser location, green square indicates ROI. B. Time course of laser-mediated VISp response as a function of laser power (solid line, mean; shaded region, S.E.M. across trials; n=35 trials per laser power from example session). Red bar indicates laser on. C. Time course of VISp response to external full-field checkerboard stimulus as a function of stimulus contrast (n=25 trials per contrast). Grey bar indicates visual stimulus present. D. VISp response across laser powers when targeting frontal cortex (light purple), somatosensory cortex (medium purple), or visual cortex (dark purple; error bars indicate S.E.M. across trials, n=90 trials across n=3 subjects). E.Schematic of the visual stimulus for retinal adaptation. White squares are shown on a black background, switching position randomly between 45-450 ms. Asterisks indicate significant visual response (paired t-test between dF/F 200 ms before and after stimulus onset across all trials, p<0.05). F. Time course of laser-mediated VISp response with and without retinal adaptation (example session, 0.9 mW laser stimulus, n=35 trials per condition). G. Magnitude of laser-mediated VISp response across laser powers with and without adaptation (n=90 trials per condition across n=3 subjects). Dashed line indicates response to full-field checkerboard stimulus of 12.5% contrast. Asterisks indicate significant visual response (paired t-test between dF/F 200 ms before and after stimulus onset across all trials, p<0.05). H. Difference in peak activity in VISp across laser powers with retinal adaptation versus without (n=90 trials per condition across n=3 subjects). Asterisks indicate significant difference (paired t-test between dF/F 200 ms before and after stimulus onset across all trials, p<0.05). I. Example calcium activity image when stimulating frontal cortex in opsin subject (left, n=45 trials, 0.4 mW laser power) or without opsin (right, n=35 trials, 0.4 mW laser power). J. Peak VISp response in control subjects (with adaptation, green, without adaptation, orange; n=90 trials per condition across n=3 subjects) and peak frontal cortex response in opsin subjects (pink, n=125 trials per condition across n=3 subjects). Asterisks indicate significant change from baseline (for opsin subjects, paired t-test between dF/F 200 ms before and after stimulus onset across n=90 trials, for control subjects, paired t-test between dF/F 300 ms before and 120 ms after stimulus onset across n=125 trials, p<0.05).

We reasoned that retinal adaptation could reduce unintended visual responses from laser stimulation. To test this, we measured the visual response to laser stimulation with and without the presence of a sparse noise visual stimulus (see Methods, Figure 3E). On trials with retinal adaptation, no significant visual response was detected at any laser power (Figure 3G, Supplemental Figure 3C). Our results indicate that inducing retinal adaptation extends the effective range of optogenetic stimulation by 0.7 mW (Figure 3J).

## Discussion

Here we developed a system for simultaneous mesoscopic measurement and manipulation of mouse cortical activity in combination with large-scale electrophysiology. We demonstrate reliable inactivation of neural populations across the dorsal cortical surface in awake mice, with these effects observable in both the widefield calcium imaging and electrophysiological recordings. The spectra of the two modalities were effectively separable: the blue and violet lights used for fluorescence imaging did not activate the opsin, and the red light used to drive the opsin did not interfere with the green imaging channel. The inactivation magnitude depended on both laser intensity and laser stimulus duration; higher laser intensities caused larger spatial inactivation across the cortex. The viral strategy remained stable over many weeks within mice, and though the magnitude of inactivation by laser power varied across subjects, the slope of the relationship between laser power and inactivation magnitude was similar. During stimulation of the frontal cortex, we observed off-target responses in the visual cortex that were reduced with a masking visual stimulus.

The spatial spread of inactivation that we observed was unexpectedly large, with significant suppression detected several millimeters from the laser target (Figure 2C) despite the small size of the focused laser beam (0.12 mm diameter, Supplemental Figure 1C). We considered that light scattering might compromise spatial precision, however, our electrophysiological observations suggest this is not the case: neurons activated by the laser (i.e. Chrimson-expressing inhibitory interneurons, which do not express GCaMP) were only activated near the laser location, though laser stimulation further away did suppress many of the other recorded neurons (presumably mostly those expressing GCaMP under the CaMKII driver; Figure 2E). The local effect of the laser on inhibitory interneurons but longer-range suppression of excitatory neurons, as reported previously with similar optogenetic strategies (Li et al., 2019), suggests that the range of suppression is determined by network propagation of the suppressive effect. In the future, our method could be used to carefully map the zone of influence of each cortical location, i.e. the specific set of areas affected by its inactivation, which we do not expect to be symmetrical.

The off-target activation of visual cortex upon stimulation of frontal cortex may pose a challenge for experiments that rely on visual stimuli or vision-based behavioral tasks. These effects almost certainly arise from direct activation of the retina by the laser from behind. Though the laser is red (638 nm) and rodents exhibit minimal sensitivity to red light relative to shorter wavelengths (Jacobs et al., 1991), behavioral responses to red light have been noted (Nikbakht and Diamond, 2021). This interpretation is supported by several observations: 1) The effect is observed in control subjects with no opsin expression, ruling out any network response to frontal cortical activity changes; 2) The part of visual cortex that becomes active (at the anterior end of VISp, Figure 3A), represents the lower visual field, i.e. the dorsal part of the retina, where light would be expected to arrive after passing through frontal cortex; 3) The effect is reduced by adapting the visual system with high-contrast stimuli, an approach previously employed for this problem (Danskin et al., 2015). Nevertheless, whether using the adapting visual stimulus or not, a range of powers that modulate frontal cortical activity but do not drive detectable visual responses is available.

This system is constrained to head-fixed mice, rather than freely moving and naturally behaving subjects. Though head-mounted widefield imaging microscopes have been developed (Scott et al., 2018; Rynes et al., 2021), additional modifications of those systems would be necessary to incorporate targetable, precise stimulation for optogenetic control. Furthermore, combining electrophysiological measurements, as we have demonstrated here, may be challenging. The present system is compatible with the large range of behaviors that have been developed for head-fixed mice.

Future studies can leverage this technique to further investigate neural correlates of behavior. Any combination of promoters may be used to drive the GCaMP sensor and Chrimson opsin expression, facilitating studies of cell type function in distributed neural circuits. The addition of high-density electrophysiology adds additional resolution and allows for linking the activity of deeper brain structures to causal perturbations and mesoscale dynamics in cortex. The ability to record and manipulate across the entire dorsal cortical surface also provides the opportunity to perform closed-loop perturbations at the mesoscale. This system will be instrumental for future research on inter-area communication during cognition, perception, and action.

## Methods

### Animals

All experimental protocols were conducted according to US National Institutes of Health guidelines for animal research and approved by the Institutional Animal Care and Use Committee at the University of Washington.

### Mouse Lines

We used double-transgenic mice obtained by crossing CaMKII-Tta (B6.Cg-Tg(Camk2a-tTA)1Mmay/DboJ; Jax #007004) with either tetO-GCaMP6s (B6;DBA-Tg(tetO-GCaMP6s)2Niell/J; Jax #024742) or tetO-jGCaMP8s mice (B6;D2-Tg(tetO-GCaMP8s)1Genie/J; Jax #037717, provided by M. Pachitariu; (Wang et al., 2023)). To express the opsin, we retro-orbitally injected mice aged P28-42 with the DLX2.0-ChrimsonR-tdTomato opsin packaged in a PHP.eB serotype AAV. The ChrimsonR particles were produced from pAAV-Syn-ChrimsonR-tdT (Addgene #59171).

In total, nine mice of both sexes were used for experiments (30 sessions total). All nine mice expressed transgenic GCaMP; eight expressed GCaMP8s and one expressed GCaMP6s. Four of eight mice expressed Chrimson under the DLX2.0 promoter (22 sessions). The other five mice had no viral injection and expressed GCaMP only (5 sessions). Two of the four Chrimson-expressing mice were used for experiments with simultaneous widefield imaging, optogenetic activation, and electrophysiological recordings (3 sessions).

### Surgery

For opsin expression, we retro-orbitally injected 4-6 week old mice with the AAV-PHP.eB-DLX2.0-Chrimson virus (typically 1.6e13 GC diluted in 100 uL PBS, see Table 1). Animals were anesthetized with isoflurane (4% in O2) and injected with virus diluted into 100-150 uL of PBS. After mice reached P48 or later, an implant surgery was performed under isoflurane (1-4% in O2). Briefly, animals were given carprofen (5 mg/kg) subcutaneously in the shoulders and lidocaine (2 mg/kg) subcutaneously in the surgical area above the skull. The skin was cleared to reveal the surface of the dorsal skull. The edges of the incision were glued to the skull with cyanoacrylate (VetBond, World Precision Instruments) to protect the underlying muscle. The skull was treated with 10% citric acid (Dentin Activator Liquid, Parkell) to improve the bond between skull and implant. Then, a 3D-printed recording chamber was secured to the skull with dental cement (Metabond, Parkell). Thin layers of

UV-cured optical glue (Norland Optical Adhesive 81, Norland Products) were applied to the exposed surface of the skull inside the chamber. A 3D-printed titanium headpost (ProtoLabs) was then cemented to the posterior end of the recording chamber. After surgery, mice were treated with carprofen for two days and given at least one week to recover before experiments.

### Microscope design

We designed the microscope to have a single shared light path to the brain, with a triple-dichroic setup. The GCaMP excitation light path involved blue (470 nm) and violet (405 nm) LEDs combined with a 425 nm dichroic mirror and coupled into a liquid-core light guide (Cairn OptoLED system). The blue/violet excitation light was introduced to the central light path at a right angle and reflected off a 495 nm dichroic mirror downward towards the 0.63x objective lens and brain. The green GCaMP emission light passed through the 495 nm dichroic mirror and reflected off a 580 nm dichroic mirror to exit the central light path at a right angle towards the 1x condenser lens and camera. The red light for optogenetic excitation was produced by a 638 nm diode laser coupled to a Ø50 μm, 0.22 NA fiber patch cable. The fiber output was collimated and coupled into a two-dimensional galvanometer mirror system to steer the beam downwards from the top of the central light path. This light passed through scan and tube lenses with long focal lengths such that the red light could travel through the long central chamber housing the 580 and 495 nm dichroics before being focused on the brain by the objective lens. See Table 2 for a complete parts list.

An alternative strategy to introduce the red laser light could have been by a separate light path with a separate objective lens, for example angled toward the brain from above and to the side of the subject. This system would have been simpler in the sense of avoiding the need to develop a way to couple the red light into the same light path as the imaging. However, our approach has several advantages: 1) There are no extra parts around the microscope relative to a standalone widefield microscope, preserving as much space as possible for combining the imaging and stimulation with electrophysiology; 2) The laser light naturally has access to the same field of view as the microscope and does not require a perspective transformation to map galvanometer voltages to coordinates on the surface of the brain. 3) The surface of the brain is as close to the focal plane of the laser as possible - if the light were introduced at a tilt from the side, the laser would be differently in focus at different points on the skull.

### Widefield calcium imaging and optogenetic experiments

For microscope part numbers and optics details, see above. Images were captured at 70 Hz and 560 × 560 pixel frame size (after 3×3 binning). The microscope uses a 0.63x objective lens with a 1x condenser lens at the camera; the resulting field of view is 9.7 × 9.7 mm. For imaging the calcium indicator, frames were alternately illuminated with blue (470 nm) and violet (405 nm) light for hemodynamic correction (see below). In order to minimize light artifacts from LED illumination during electrophysiology, the onset and offset of all LED pulses were ramped over 1 ms in a sinusoidal shape. For optogenetic activation, a 638 nm laser was fixed above the microscope scan and tube lenses. The position of the laser was controlled by two mirror galvanometers. During experiments, mice were headfixed under the objective and behind three 60 Hz refresh rate screens (Adafruit LG LP097QX1). A 3D printed plastic cone glued to light-blocking fabric (Thorlabs BK5) was fixed to the implant well and screens to prevent light contamination. The mouse forepaws could turn a rotating rubber wheel with a rotary encoder. A camera (FLIR Chameleon3) with IR filter and 16 mm focal length lens (Thorlabs MVL16M23) captured face and paw movements.

At least two days before experiments, mice were acclimated to handling and head fixation. In order to maintain an awake and alert state, seven out of nine mice were lightly water-restricted for at least one day before beginning imaging sessions (three out of four of the opsin-expressing mice and four of the five GCaMP-only mice, Table 1). These mice received 10% sucrose rewards in random 2-5 second intervals throughout experiments.

During experiments with optogenetic stimuli only, mice were also shown sparse noise stimuli on the screens. The stimulus consists of a black background with six white blocks, which randomly move in position every 45-450 ms. The sparse noise was also used for experiments testing the effect of visual responses to laser stimuli (Figure 3E-H).

Experiments in Figures 1 and 3 used a 100 ms step function laser stimulus. The onset and offset of the laser ramped in power in a sinusoidal shape over 1 ms. For experiments with longer stimulus durations using a sine wave laser stimulus (Figure 2), the waveform was adjusted such that the trough was zero and the peak was twice the target power (such that the mean power over time matched the target power). In order to avoid unintended calcium activity from the sound of the galvanometers moving, we added a uniformly random 100-800 ms delay between the movement of the galvanometers and the laser stimulus onset. Laser power and galvo position were controlled via analog output from a NI DAQ (National Instruments USB6343).

In order to measure laser power, we used a light meter (Thorlabs PM100D with S130C) to record power in milliwatts. We made 14 measurements in the voltage modulation range (0-5 V) of the laser. The relationship between input voltage and measured power was linear, and consistent across sessions. To compute laser intensity (power per area), we measured the beam diameter using a mirror positioned at the sample plane. The beam power, when reflected by this mirror, is high enough that the small portion reflected by the 550 nm longpass dichroic is detectable on the camera. During *in vivo* imaging, the skull preparation does not reflect enough laser power to be detected.

We computed the LED intensity for blue and violet imaging using the area of the photodiode sensor (Thorlabs S130C), which has a diameter of 9.5 mm (area of 70.8 mm^2^).

### Widefield data post-processing and analysis

Non-neuronal signals arising from hemodynamic changes were removed by subtracting the violet-light evoked signals from the blue-light evoked signals with linear regression, as previously described in (Zatka-Haas et al., 2021; Ye et al., 2023). To store and process the data, we compressed the widefield data

1. into spatial components (U) and temporal components (S*V) with singular value decomposition in the form D = USV^T^. We used the top 50 singular vectors, which accounted for 98.2 ± 0.2% (mean ± SEM, 9 subjects) of total variance before compression. In analyses describing calcium activity timecourses, we averaged across a 1.7 × 1.7 mm (100 × 100 pixels) region of interest around the laser target. To compute stimulus-aligned dF/F, we used a baseline of either the mean image or the mean of four frames (114 ms) before stimulus onset.

We computed the duration of significant inhibition (as in Figure 2A, B) via a threshold determined from a null distribution. The null distribution consisted of calcium activity at baseline (300 ms before stimulus onset), and the threshold was mean-2*S.D of the distribution. To determine the spatial extent of activation (as in Figure 2C, D), we similarly computed a null distribution from baseline (150 ms before stimulus onset), and thresholded (mean-2*S.D.) all pixels. We then converted pixels to millimeters based on measurements of the microscope field of view (Supplemental Figure 1A, 17.3 μm/pixel).

For visual response experiments (Figure 3), we averaged across a 519 × 519 μm (30 × 30 pixels) region of interest to match the area of the laser-evoked visual response.

### Electrophysiology experiments

Recordings were made with four-shank Neuropixels 2.0 probes (Steinmetz et al., 2021). On or before the first day of recording, a 2-3 mm diameter craniotomy was prepared over left VISp under anesthesia. The center of the craniotomy was 2.5 mm left of lambda. The craniotomy was protected with transparent dura-gel (Dow Corning 3-4680 Silicone Gel), with a 3-D printed protective cap secured above the implant well. After at least two hours of recovery, mice were head-fixed in the electrophysiology rig. We used internal referencing for all recordings, inserting the probe directly through the dura-gel without a saline bath. The probe was inserted at a 50° angle to accommodate the working range of the objective (7.5 cm). We used SpikeGLX for data collection and Kilosort 4 (Pachitariu et al., 2024) via SpikeInterface (Buccino et al., 2020) for spike sorting. Although light artifacts resulting from laser illumination can be minimized with a ramped stimulus (see above), we ensured artifact removal by excluding spike times within 1 ms of laser onset and offset. Spikes were binned at 10 ms. Units were excluded from analysis if their overall spike rate was below 0.1 spikes/second (approximately 60% of units excluded).

## Acknowledgements

We thank Marius Pachitariu for contributing the tetO-jGCaMP8s mouse line. We thank Michael Krumin for consultation on microscope design. We thank Tanya Daigle for consultation on the genetic strategy. We thank Ljuvica Kolich for assistance with mouse husbandry and breeding. We thank the Allen Institute for Brain Science Viral Technology team for viral packaging and titer determination. This work was supported by the NIH National Eye Institute (T32EY07031 to AJL, 1F31EY035880 to AJL), the Simons Foundation Shenoy Undergraduate Research Fellowship (PM), the Pew Biomedical Scholars Program (NAS), the Klingenstein-Simons Fellowship in Neuroscience (NAS), the National Science Foundation (award 2142911 to NAS and award 2024364 to NAS), and the NIH BRAIN Initiative (UG3MH120095 to JTT).

## Author contributions

NAS designed and built the microscope. NAS designed the genetic strategy in consultation with JTT. XO-A cloned the construct for the opsin virus. ZY performed retroorbital virus injections. AJL, ZY, and AJB performed surgeries. PM and AJL collected the data, with additional contributions by ZY and AJB. PM and AJL performed data analysis and visualization. PM, AJL, and NAS wrote the first draft, and all authors reviewed and edited. NAS supervised the work.

## Supplemental Figures

**Supplemental Figure 1.**
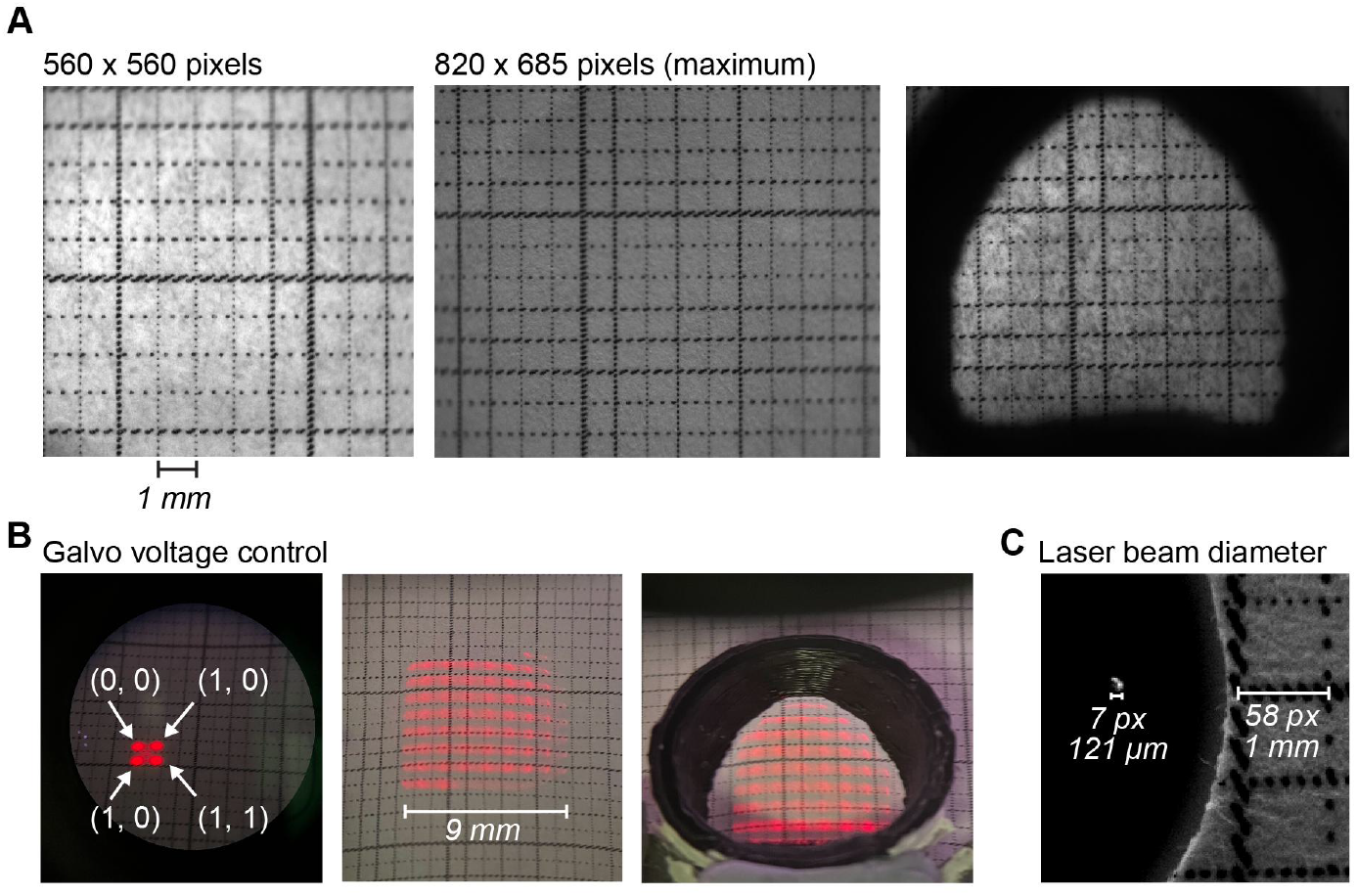
A. Frames captured by widefield microscope with ruler placed in the field of view. Each square is 1 × 1 mm. Left, 560 × 560 px FOV (after 3×3 binning) as used in experiments. Middle, maximum FOV size after 3×3 binning; camera sensor is 2464 × 2056. Right, field of view with implant well. B. Distance of galvo-directed laser movement with analog voltage control. Each square is 1 × 1 mm. Left, coordinates indicate (X, Y) galvo position in volts. Middle, working range of galvo-directed laser movement. X and Y galvo targets span -2 to 5 V in 1 V steps. Right, range of movement with implant well. C. Diameter of the laser beam via mirror placed at sample plane. Each pixel is 17.3 µm.

**Supplemental Figure 2.**
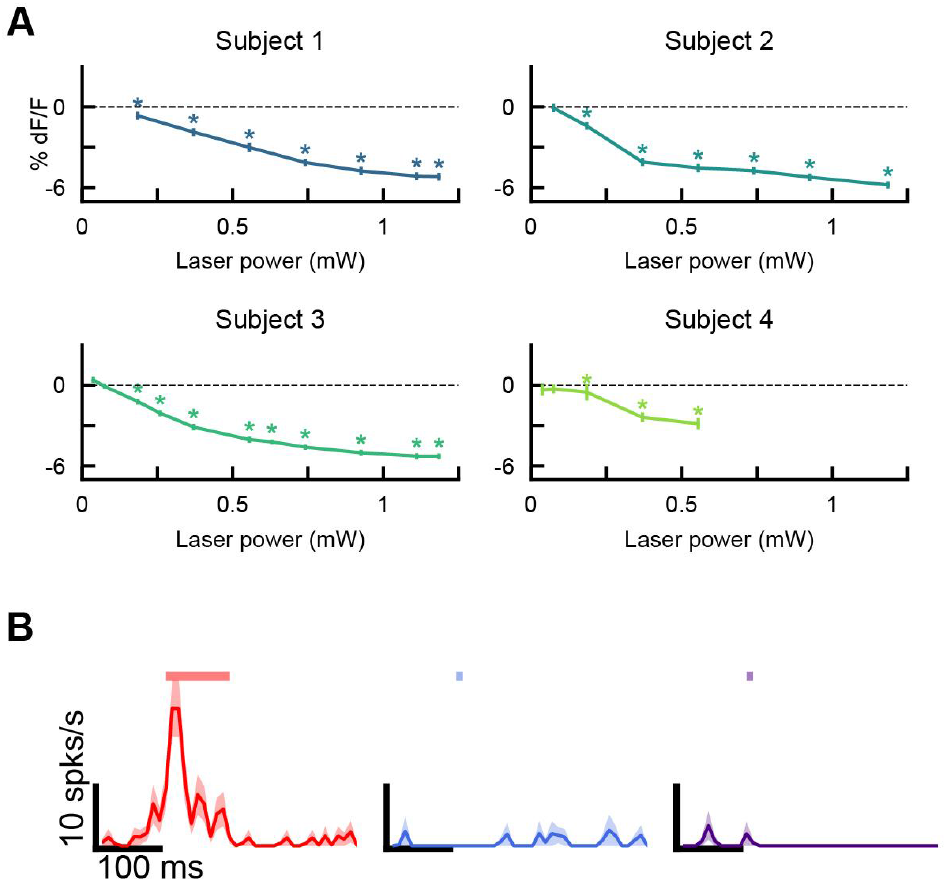
A. dF/F response from one pixel as a function of laser power (solid line, mean; error bars, S.E.M. across trials; n=40, 45, 45, 45 trials per subject). Subjects labeled as in Table 1 and Figure 2. Asterisks indicate significant response to laser power condition (paired t-test between spike counts 100 ms before and after laser onset, p<0.05). B. Example opsin-expressing neuron recorded with electrophysiology. Left, response to 0.7 mW laser stimulus (n=30 trials). Middle, response to blue LED pulse (n=40 trials). Right, response to violet LED pulse (n=40 trials). Duration (11 ms) and intensity of LED pulses are matched to experiment conditions.

**Supplemental Figure 3.**
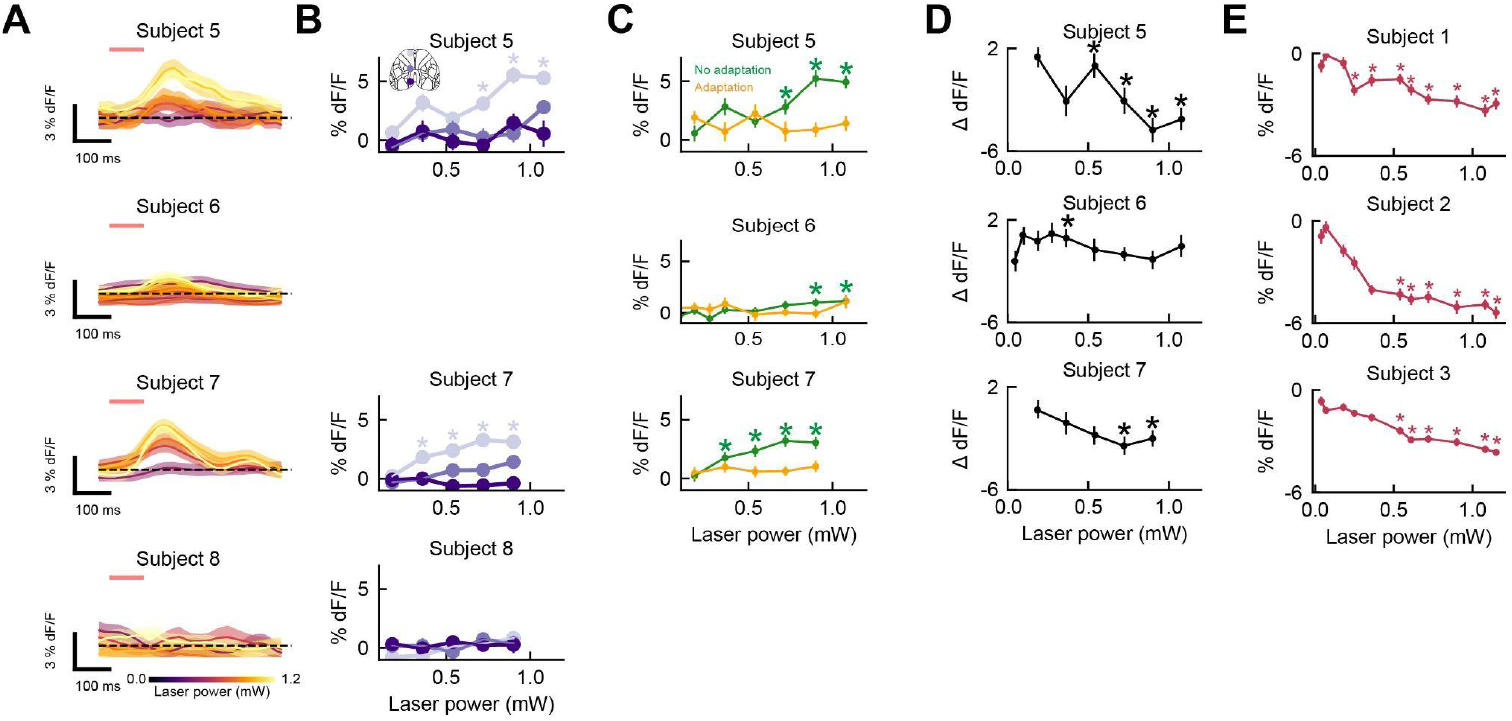
A. Timecourse of VISp response to laser stimulation in frontal cortex in control subjects (subjects as labeled in Table 1; solid line, mean; shaded regions, S.E.M across trials; top to bottom: n=28, 45, 35, 35 trials). B. VISp response to laser stimulation of frontal, somatosensory, or visual cortex (30×30 pixel ROI; error bars, S.E.M across trials; top to bottom: n=28, 35, 35 trials). Asterisks indicate significant visual response to laser power (paired t-test between dF/F 200 ms before and after laser onset, p<0.05) C. VISp response to frontal cortex laser stimulation with (orange) or without (green) retinal adaptation across subjects (top to bottom: n=28, 45, 35 trials). Asterisks indicate significant visual response to laser power condition (paired t-test between dF/F 200 ms before and after laser onset, p<0.05). D. Difference in laser-mediated VISp response with and without adaptation (top to bottom: n=28, 45, 35 trials). Asterisks indicate a significant difference (paired t-test between dF/F 200 ms after laser onset with and without adaptation). E. dF/F response in frontal cortex to laser targeted in frontal cortex in subjects with the opsin as labeled in Table 1 (top to bottom: n=40, 45, 45 trials). Asterisks indicate significant response to laser power (paired t-test between dF/F 300 ms before and 150 ms after laser onset, p<0.05).

